# CFMF: A Clustering-Free Cell Marker Finder for Single-Cell Transcriptomics Data

**DOI:** 10.1101/2025.10.26.683836

**Authors:** Shumin Yin, Ke Liu

## Abstract

**Background:** Accurate identification of marker genes is essential in single-cell RNA sequencing (scRNA-seq) analysis. However, conventional clustering-based annotation is constrained by resolution parameter selection and predefined marker gene databases.

**Methods:** We developed Clustering-Free Cell Marker Finder (CFMF), a novel computational framework that enables the discovery of marker genes without clustering. CFMF was validated using multiple scRNA-seq datasets and systematically benchmarked against widely used gene selection methods.

**Results:** We first validated CFMF using the PBMC3K dataset and found that it not only recovered canonical markers, but also uncovered novel marker genes. When validating with glioblastoma datasets, CFMF successfully recapitulated established subtype signatures. Using scRNA-seq data of human normal lung, we demonstrated its superior sensitivity in rare cell detection even at very low prevalence. We applied CFMF to perform integrative analysis on colorectal cancer scRNA-seq data and defined seven transcriptionally distinct subclones, which includes an *LGR5*^+^ population that is strongly associated with metastasis.

**Conclusion:** CFMF represents a versatile and robust strategy for marker gene discovery, rare cell detection, and integrative dissection of tumor heterogeneity, thereby advancing single-cell research and facilitating the identification of previously unrecognized cell types.

## Introduction

Recent advances in single-cell transcriptomic technologies have enabled the construction of comprehensive cellular atlases across diverse species and organs[1-5], providing novel insights into tissue architecture and cellular function. A critical step in scRNA-seq data analysis is cell type annotation, which involves assigning individual cells to established cell types or functional states[6]. Annotation strategies can be broadly categorized into manual and automated approaches: manual annotation defines cell identity primarily based on the expression of known marker genes within cell clusters, whereas automated annotation typically leverages machine learning or statistical modeling methods to infer cell identity by comparing single-cell transcriptomic profiles with reference datasets representing established cell types[7].

Despite its widespread adoption, manual annotation exhibits several limitations. First, it inherently depends on the availability of highly specific marker genes[8] and requires extensive domain knowledge, thereby introducing subjectivity into the annotation process. This limitation is particularly evident in the tumor subtyping, where substantial intratumoral heterogeneity[9, 10] obscures the delineation of distinct subclonal populations and hinders the identification of reliable subtype-specific markers. Second, manual annotation typically begins with clustering, yet the clustering outcome is strongly influenced by parameter choices (e.g., the number of clusters). The use of inappropriate parameters may artificially split a single cell type into multiple clusters or fail to capture rare populations[11]. In particular, rare cell types[12, 13] are often masked by more abundant populations, leading to their misclassification and the missed opportunity to discover novel cell types. Therefore, there is an urgent need to develop clustering-independent methods that can robustly identify highly specific marker genes and thereby improve the reliability of cell type annotation.

In this study, we present CFMF, an innovative framework for marker gene identification that operates independently of both clustering and prior domain knowledge. CFMF is founded on the hypothesis that cells expressing the same marker gene exhibit spatial proximity in the transcriptomic landscape, thereby forming localized aggregations. We evaluated CFMF across multiple normal and malignant datasets and demonstrated its ability to recover both canonical markers of established cell types and novel markers, including those specific to rare populations. Moreover, when using CFMF to delineate intratumor heterogeneity of colorectal cancer (CRC), we uncovered seven transcriptionally distinct CRC subclones. Notably, subclone 4 was enriched in metastatic CRC, and exhibited an inverse correlation with CD8^+^ T cell infiltration, implicating a potential role in immune evasion and metastatic progression. Finally, we systematically compared CFMF with other established gene selection methods and found that CFMF identified more biologically meaningful genes.

## Materials and Methods

### Data acquisition and preprocessing

A total of 17 scRNA-seq datasets were collected (**Table 1**), primarily from 3CA[14], the Single Cell Portal (https://singlecell.broadinstitute.org/single_cell), and TISCH2[15]. Raw gene expression matrices were processed using Seurat (v4.3.0)[16]. Dimensionality reduction was performed using principal component analysis (PCA) and Uniform Manifold Approximation and Projection (UMAP) was computed on the first 30 principal components using Seurat default parameters for two-dimensional visualization.

### Cell type annotation

SingleR was used to perform automatic cell type annotation with the BlueprintEncodeData as the reference[17]. Copy number variation (CNV) profiles were inferred by inferCNV (v1.12.0) (using immune cells as the reference population)[18], and were further used to identify malignant cells.

### The full CFMF pipeline

The CFMF pipeline comprised the following steps:

1. Gene Expression Filtering. Genes detected in at least 3 cells but not expressed in all cells were retained.
2. Dimensionality reduction. PCA was performed on HVGs and PCs were used for subsequent neighbor calculations.
3. Local neighborhood enrichment. For each gene, positive cells were defined as those with nonzero expression. For each positive cell, the 50 nearest neighbors were identified in PCA space using get.knn in the FNN package with Euclidean distance. Fisher’s exact test was applied per positive cell to assess enrichment of positive neighbors, and per-cell *p*-values were combined into a single meta *p*-value using meta-analysis.
4. Global compactness. Silhouette[19] values were computed in PCA space using binary labels (positive vs non-positive) for each positive cell; the median silhouette across positive cells was recorded.
5. Threshold selection and validation. Thresholds of -log_10_(meta *p*) (100, 200, 300, 400) and silhouette (-10, 0, 10, 20) were evaluated on the PBMC3K dataset[20]. Operational cutoffs of -log10(meta *p*) > 200 and silhouette > 0 were selected, where CFMF-selected genes yielded consistent recovery of canonical markers. Known marker genes were curated from CellMarker 2.0[8].

### Pathway activity inference

Single-cell pathway activity was computed with AUCell (v1.20.2)[21] on raw RNA counts using MsigDB hallmark[22] and Reactome[23] gene sets. The active region was defined as the top 5% of expressed genes per cell (aucMaxRank = 5% of all genes). Hallmark and Reactome gene sets were obtained from the Enrichr database[24].

### Immune infiltration analysis

For each CRC scRNA-seq sample, the proportion of immune cells within the non-malignant compartment, and the relative abundance of each CRC subtype among malignant cells were calculated. Associations between the two values were assessed using the Spearman rank correlation.

### Site distribution preference of CRC subclones

*R*_*o/e*_ (Ratio of observed to expected frequency) was calculated with the Startrac (v0.1.0) package[25]. The R_*o/e*_ value was defined as:

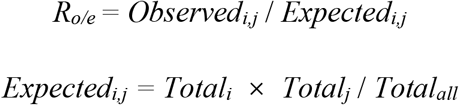

where *Observed*_*i,j*_ is the number of cells from subclone *i* in tissue *j, Total*_*i*_ is the total number of cells in subclone *i, Total*_*j*_ is the total number of cells in tissue *j*, and *Total*_*all*_ is the total number of all cells across both tissues.

### Spatial transcriptomic data analysis

A total of 26 CRC metastasis tumor sections were curated for analysis. ESTIMATE scores were computed per spatial spot using the ESTIMATE R package (v1.0.13), and converted to tumor purity via the established formula[26]:

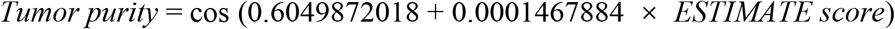

For each section, k-means clustering (k = 5) was applied to normalized expression profiles and the two clusters with highest ESTIMATE scores were designated as neoplastic.

The spatial deconvolution was performed with spacexr package (v2.2.1)[27]. A reference matrix was constructed by random sampling 5,000 malignant CRC cells; subclone proportions per spatial spot were estimated using the RCTD algorithm implemented in spacexr.

### Comparison with established feature-selection methods

Feature selection methods (Fano[28], HVG[16], M3Drop[29], RaceID3[30], SCTransform[31], and Entropy (SE)[32]) were applied to derive method-specific gene sets. For each gene nominated by a given method, cell-type specificity was quantified by the area under the curve (AUC) produced by the scoreMarkers function in scran package[33]. The largest min.AUC across all cell types was regarded as the final AUC.

## Results

### I. Overview of the CFMF method

To address the limitations of conventional cell type annotation, which relies heavily on clustering algorithms and predefined marker sets, we developed a cluster-independent algorithm termed CFMF. CFMF is founded on the hypothesis that cells expressing the same marker gene exhibit inherent transcriptional similarity and thus localize adjacently in transcriptomic space. Guided by this principle, CFMF first applies PCA to reduce dimensionality of the expression matrix, and subsequently designates a gene as a candidate marker if it demonstrates consistent expression across spatially adjacent cells while remaining largely absent in the remaining (**Figure 1A**).

Given a gene *g*, we first identified all cells expressing *g* (termed as *g*^+^ cells). Specifically, for each *g*^+^ cell, we then quantified the local enrichment of *g*^+^ cells by applying Fisher’s exact test to its 50 nearest neighbors in PCA space (**Figure 1A**). The resulting *p*-values derived for each *g*^+^ cell were subsequently integrated via meta-analysis, yielding a single meta *p*-value that reflects the overall degree of local enrichment for gene *g*. Recognizing that local metrics alone may fail to capture global distribution patterns, we further computed silhouette coefficients of *g*^+^ cells to assess their transcriptomic cohesiveness (**Figure 1A**). Accordingly, an ideal marker gene is expected to exhibit both a low *p*-value and a high silhouette coefficient. Thresholds for both the *p*-value and silhouette score were established through pilot experiments (**Methods**), with genes that satisfy the criteria of -log_10_(meta *p*-value) > 200 and silhouette score > 0 being classified as potential markers in the scRNA-seq datasets.

### II. Validating CFMF with public scRNA-seq datasets

To validate the performance of CFMF, we first applied it to the PBMC3K dataset, which comprises 2,700 peripheral blood mononuclear cells from 10x Genomics[20]. As expected, CFMF accurately recovered canonical lineage markers of mononuclear cells, including *S100A8* for myeloid cells, *CD3E* for CD4^+^ T cells, and *CD14* for classical monocytes (**Figure 2A, Figure S1A**). In addition, CFMF also identified subtype-specific markers, such as *CCR7* and *LEF1*, which specifically expressed in CD4^+^ naïve T cells (**Figure 2A, Figure S1A**). Notably, CFMF also uncovered several novel marker genes for known cell types. For example, *PRSS23* and *AKR1C3* were broadly expressed in NK cells, *FOLR3* and *RBP7* in CD14^+^ monocytes, *HMOX1* in CD16^+^ monocytes (**Figure 2A-2B, Figure S1B**), thereby demonstrating the ability of CFMF to identify cell type–specific markers.

To assess the generalizability of CFMF, we next applied it to a glioblastoma (GBM) scRNA-seq dataset[34] comprising five molecular subclones. As anticipated, the CFMF-selected genes recapitulated previously reported GBM subclone signature genes[34], including *CST3* and *S100B* for the astrocytic (AC) signature, as well as *BCAN* and *GPR17* for the oligodendrocyte progenitor cell (OPC) signature (**Figure 2C**). Moreover, CFMF revealed additional genes not included in previous signature lists but displaying pronounced subclone-specific expression, including *LGALS3* and *TNFRSF1A* for the mesenchymal-like (MES) signature, *MLC1* and *AQP4* for the AC signature, *TNR* and *COL20A1* for the OPC signature, and *DCX* and *ELAVL4* for the neural progenitor-like (NPC) signature (**Figure 2C-2D, Figure S1C**). These results demonstrate that CFMF effectively reveals the transcriptional heterogeneity of GBM.

Furthermore, we assessed the ability of CFMF-selected genes to recover known markers in both the PBMC3K and GBM datasets. Interestingly, CFMF-selected genes were significantly enriched for upregulated DEGs across multiple cell types in PBMC3K, as well as for established GBM subclone signature genes, when compared with non-selected genes (OR = 73.7, *p* = 1.65 × 10^-65^, and OR = 50.4, *p* = 1.18 × 10^-166^, respectively; Fisher’s exact test; **Figure 2E-F**). Collectively, CFMF provides a robust framework for identifying cellular heterogeneity across both normal and malignant populations.

### III. CFMF fuels the identification of rare cell types

Growing evidence suggests that rare cell types are pivotal in development, immune responses, and disease progression[35, 36]. Yet, their scarcity across tissues renders them difficult to resolve by conventional clustering-based scRNA-seq methods, particularly when confined to individual samples. Given the demonstrated ability of CFMF to identify biologically informative marker genes, we investigated whether it could also be applied to detect rare cell types. To this end, we first curated several normal lung scRNA-seq datasets previously reported to contain rare populations[3], such as ionocytes and pulmonary neuroendocrine cells, and applied CFMF on each individual sample level to identify potential marker genes. Among the 42 samples subjected to individual clustering analysis, ionocytes did not form a distinct Seurat cluster in 28 cases, prompting us to focus subsequent analyses on these samples. When examining genes expressed in fewer than 5% of cells, we found that CFMF successfully identified markers of rare cell types, such as *SCG3* for pulmonary neuroendocrine cells (**Figure 3A, Figure S2A-S2D**), and *ASCL3* and *BSND* for ionocytes (**Figure 3B, Figure S2A-S2D**). Remarkably, CFMF was able to recover ionocyte markers even when their prevalence within a sample declined to as low as 0.02%. Across samples containing more than three cells of these rare populations, 88% exhibited detectable marker genes identified by CFMF. Collectively, these findings highlight the superior sensitivity of CFMF in detecting transcriptional signatures of rare cellular populations.

### IV. CFMF as a powerful tool for delineating tumor heterogeneity

Tumor cell heterogeneity constitutes a critical barrier to effective therapeutic intervention, underscoring the necessity of delineating its molecular determinants[9, 10, 37]. Building on our previous findings that CFMF can effectively identify subclone-associated markers in GBM, we applied CFMF to the scRNA-seq datasets of 109 CRC samples to identify CRC subclones. As the identified genes might include markers of non-malignant cells (such as *NKG7* for NK cells), we implemented additional filtering steps to refine the CFMF-selected candidates (**Figure 4A**). For each sample, genes were processed as follows: (i) genes meeting the CFMF criterion were identified; (ii) genes for which more than half of the expressing cells are malignant were retained; (iii) genes expressed in more than half of all malignant cells were excluded; (iv) the top 1,000 genes ranked by silhouette were selected. The candidate gene sets from all samples were then aggregated, and the 2,000 most recurrent genes were designated as the final marker set. Notably, substituting these CFMF-selected genes for HVGs during batch correction enabled effective integration of malignant cells across samples, successfully mitigating batch effects that conventional HVG-based approaches failed to resolve (**Figure 4B**). These findings indicate that the recurrent CFMF-selected genes capture core biological features that are shared across diverse cancer samples.

Clustering analysis of malignant cells was performed using the top 2,000 recurrent genes. After excluding 6 low-quality clusters characterized by reduced transcript counts and elevated ribosomal gene expression (**Figure S3A**), the remaining cells were reclustered, yielding 8 distinct subclones (**Figure 4C**). To characterize the biological states of these subclones, we quantified MSigDB hallmark gene sets activities[22] in each malignant cell using AUCell[21] and compared the activity profiles across subclones (**Figure 4D**). Subclone 1, marked by *S100A11* and *PHLDA2*, is predominantly up-regulated epithelial-mesenchymal transition and inflammatory responses pathways, while subclone 2, defined by *ADH1C* and *OLFM4*, exhibited dysregulation of metabolic pathway. Subclone 3 demonstrated a hybrid profile, combining features of both subclone 1 and subclone 2, thereby suggesting an intermediate state. Subclone 4 displayed upregulation of the WNT signaling pathway and subclone 5 is distinguished by the activation of MYC pathway, while subclone 6 lacked dominant hallmark pathway activities. Subclone 7 and subclone 8 share highly similar MYC pathway upregulation and proliferative characteristics, prompting their consolidation into a single subclone. The robustness of these subclone-specific characteristics was further validated by Reactome pathway analysis[23] (**Figure S3B**), which corroborated the hallmark-derived profiles. Consequently, we identified a total of 7 subclones with distinct biological characteristic (**Figure S4A-S4B**).

When comparing subclonal distributions between primary tumors and metastatic sites, we observed that primary lesions were predominantly composed of subclones 2, 3, 6 and 7, whereas metastatic sites were dominated by subclones 1, 4 and 5 (**Figure 4E**). Notably, subclone 4 showed the most pronounced enrichment at metastatic sites (**Figure 4E**) and was therefore prioritized for further analysis. This subclone was characterized by elevated expression of *LGR5* (**Figure S4B**), a gene previously implicated in metastatic progression[38, 39]. Across samples, the proportion of subclone 4 was significantly negatively correlated with CD8^+^ T cell abundance (**Figure 4F**), suggesting an immune-evasive or immune-poor state. Together, these finding underscores a potential role for subclone 4 in tumor metastasis. In addition, we compared the proportion of these subclones in malignant spots using a CRC spatial transcriptomic data[40], including 26 metastatic slices (**Methods**). The results revealed that subclone 4 predominated within metastatic regions (**Figure 4G**), further highlighting its association with metastasis.

### V. Comparison of CFMF with established methods

To further assess the ability of CFMF to identify biologically informative genes, we benchmarked it against several established gene selection methods, including Fano, HVG, M3Drop, RaceID3, SCT, and SE. We wondered whether CFMF-selected genes could more effectively capture biological distinctions among cell types by exhibiting cell type– specific expression patterns. For each method, we calculated the AUC for the identified marker genes to quantify their expression specificity across cell types in the PBMC3K dataset and 4 GBM datasets. Notably, CFMF consistently outperformed all other methods in most cases, achieving higher AUC scores across a broad range of gene selection thresholds (top 30 to 2000 genes) (**Figure 5A-B, Figure S5A-S5B, S6A-S6B**), highlighting the effectiveness of CFMF in prioritizing biological meaningful genes in both normal and malignant single-cell transcriptomic data.

Given that genes exhibiting elevated expression in specific cell types generally yield higher AUC values, we next investigated whether CFMF-selected genes enhance the discrimination across cell types. To this end, we applied two complementary validation strategies. Firstly, we assessed the clustering performance of genes identified by different methods, using silhouette coefficients[19] as the quantitative metric. Specifically, these genes were substituted for HVGs in the standard Seurat clustering workflow. Strikingly, CFMF-selected genes yielded higher silhouette scores across multiple cell types in the PBMC3K dataset compared with those obtained from alternative approaches (**Figure 5C**), indicating that CFMF-selected genes provide enhanced discrimination among distinct cell types.

As a complementary validation strategy, we applied the SingleR annotation framework[17] on the PBMC3K dataset using a repeated random sampling approach. In each of 50 iterations, 70% of the cells were randomly sampled to construct a training set, from which signature genes were identified by each method. The resulting marker genes were then used to train independent reference models, which subsequently predicted the cell identities of the remaining 30% in the validation set. Notably, across all iterations, CFMF-selected genes achieved significantly higher classification accuracy than those identified by alternative approaches (**Figure 5D**). Collectively, these results demonstrate that CFMF-selected genes encompass a larger proportion of biologically informative genes reflecting cell type-specific signatures than conventional methods, thereby facilitating more precise discrimination of distinct cellular populations.

## Discussion

In this study, we introduce CFMF, a clustering-independent computational framework designed to identify candidate marker genes from scRNA-seq datasets. The method is built on the assumption that if a gene is a marker gene of a specific cell type, then cells expressing this gene should exhibit relatively higher transcriptomic similarity. To evaluate its performance, we applied CFMF across multiple independent datasets and consistently observed robust results. For instance, in the PBMC3K dataset, CFMF recovered well-established canonical markers such as *CD14* for classical monocytes and *LEF1* for CD4^+^ naïve T cells, while also uncovering novel but highly cell type– specific markers, such as *AKR1C3* for NK cells. Similarly, in the glioblastoma dataset, CFMF successfully identified previously reported GBM subtype signature genes, including *MLC1* for AC signature, and *TNR* for OPC signature. Based on this validation, we further examined the utility of CFMF across three distinct analytical contexts, thereby highlighting its versatility and broad applicability to diverse single-cell transcriptomic studies.

In our view, one important application of CFMF is to detect rare cell types. A persistent challenge in clustering-based cell classification lies in the selection of clustering parameters. If the resolution is too low, rare cells are merged with other populations; conversely, if the resolution is set too high, a single homogeneous cell type may be erroneously fragmented into multiple clusters. Both scenarios ultimately lead to misleading interpretations. Established tools for rare cell identification, such as CellSIUS[41], RaceID[5], and GiniClust[42], largely rely on clustering or related computational strategies, and therefore remain susceptible to these parameter-dependent pitfalls. By contrast, CFMF adopts a fundamentally distinct paradigm. A notable strength of CFMF is its capacity to capture intra-cluster heterogeneity, thereby revealing the potential existence of previously unrecognized subtypes or rare cell populations. For instance, in normal lung scRNA-seq datasets, CFMF successfully identified rare cell markers such as *ASCL3* and *BSND* for ionocytes, providing molecular evidence for the presence of these rare cell types. Thus, unlike tools explicitly developed for rare cell detection, the primary utility of CFMF lies in its auxiliary role in highlighting rare or atypical populations, particularly in cases where clustering resolution is insufficient to delineate subtle heterogeneity.

Large-scale integrative analysis of scRNA-seq datasets has become increasingly prevalent in recent research, yet one of its major challenges lies in effective batch correction[43-45]. In conventional workflows for batch effect correction, an essential step is the selection of appropriate genes for dimensionality reduction. Ideally, these genes should capture genuine biological variation rather than technical variability introduced by batch effects. Since CFMF was applied in each individual sample, which mitigates the effect of batch on gene expression. Consequently, the expression variation of CFMF-selected genes is more likely to reflect underlying biology rather than technical artifacts. Although we cannot completely exclude the possibility of residual technical variation, integration leveraging recurrent CFMF-selected genes demonstrates markedly greater robustness compared with HVG-based PCA. For example, when HVGs were used for PCA followed by Harmony integration, the *LGR5*^+^ and *TOP2A*^+^ subclones were dispersed across clusters (**Figure S7A**). In contrast, substituting HVGs with CFMF-selected genes enabled robust integration and precise delineation of these biologically relevant subclones, thereby underscoring the pivotal role of gene selection in PCA-based analyses.

Taken together, these findings demonstrate that CFMF-selected markers mitigate the impact of technical variation while retaining the capacity to capture biologically relevant heterogeneity, thus providing a powerful framework for resolving tumor subclones and dissecting the complexity of cancer ecosystems.

## Supporting information

main figure 1

main figure 2

main figure 3

main figure 4

main figure 5

## Abbreviations

CFMF: Clustering-Free Cell Marker Finder;
scRNA-seq: single-cell RNA sequencing;
CRC: colorectal cancer;
PCA: principal component analysis;
DEGs: differentially expressed genes;
GBM: glioblastoma;
AC: astrocytic;
OPC: oligodendrocyte progenitor cell;
OR: Odds Ratio;
MES: mesenchymal-like;
NPC: neural progenitor-like;
HVGs: highly variable genes;
AUC: area under the curve;
CNV: copy number variation;
UMI: Unique Molecular Identifier.

## Acknowledgments

The authors would like to thank all the public data sources involved in the present study.

## Author contributions

S.Y. and K.L. conceived and designed the study. S.Y. performed formal analysis. All authors contributed to writing, reviewing, and editing the manuscript and approved the manuscript.

## Data Availability

All datasets utilized in this study are publicly available. The corresponding repositories and accession numbers are provided in Tables 1.

## Code Availability

R scripts used to analyze data and generate figures are available upon request to the corresponding author.

