## Supplementary material for "CFMF: A Clustering-Free Cell Marker Finder for Single-Cell Transcriptomics Data": legends

**Figure 1**

**Overview of the CFMF method.**

**A.** Schematic illustrating the CFMF pipeline. Individual points in PCA space represent single cells. Grey points indicate cells with no detectable expression of the indicated gene (gene-negative), whereas pink points indicate cells with detectable expression (gene-positive). Right panels show the expression patterns of an ideal marker (top, cells expressing this gene are colored blue) and a non-marker (bottom, cells expressing this gene are colored green).

**Figure 2**

**Validation of CFMF using public single-cell RNA-seq datasets.**

**A.** Scatterplot of silhouette coefficient versus meta-*p* value for all genes in the PBMC3K dataset. Points colored in blue denote candidate marker genes (−log_10_(meta-*p*) > 200 and silhouette coefficient > 0); selected genes are labeled.

**B.** UMAP plots of the PBMC3K dataset showing annotated cell types and novel markers identified by CFMF.

**C.** Scatterplot of silhouette coefficient versus meta-*p* value for all genes in a representative GBM sample. Blue points denote candidate markers (−log_10_(meta-*p*) > 200 and silhouette coefficient > 0); selected genes are labeled.

**D.** UMAP plots of the GBM dataset showing four GBM subclone signature scores (left) and the expression of the corresponding genes identified by CFMF (right).

**E.** Bar plot comparing the distribution of the top 10 upregulated DEGs per cell type in the PBMC3K dataset between CFMF-selected and non-CFMF genes. *P*-values were calculated by two-tailed Fisher's exact test.

**F.** Bar plot comparing the distribution of subclone-specific signature genes in the GBM dataset between CFMF-derived and non-CFMF genes. *P*-values were calculated by two-tailed Fisher's exact test.

**Figure 3**

**CFMF facilitates identification of rare cell types.**

**A.** Scatterplot of silhouette coefficient versus meta-*p* value for genes expressed in <5% of cells in a representative human lung sample. Blue points denote candidate markers (−log_10_(meta-*p*) > 200 and silhouette coefficient > 0); selected genes are labeled. UMAP plots showing annotation and proportion of neuroendocrine cells, expression of the CFMF-identified marker, and Seurat cluster assignments obtained with default resolution, respectively.

**B.** Scatterplot of silhouette coefficient versus meta-*p* value for genes expressed in <5% of cells in a representative human lung sample. Blue points denote candidate markers (−log_10_(meta-*p*) > 200 and silhouette coefficient > 0); selected genes are labeled. UMAP plots showing annotation and proportion of ionocytes, expression of the CFMF-identified marker, and Seurat cluster assignments obtained with default resolution, respectively.

**Figure 4**

**CFMF as a powerful tool for delineating tumor heterogeneity.**

**A**. Flow chart summarizing the pipeline used to identify and prioritize markers of malignant cells in CRC. For an identified gene *g*, malignant.pct among pos.cells denotes the proportion of malignant cells among all cells expressing gene *g*, and pos.pct among malignant.cells denotes the proportion of cells expressing gene *g* among all malignant cells. And top 1,000 genes refer to the 1,000 genes with the highest silhouette scores.

**B**. UMAP plots demonstrating the integration of 109 CRC samples using top 2,000 HVGs (left) and CFMF-selected genes (right). Different colors represent different samples.

**C**. UMAP plot of 8 distinct CRC subclones.

**D**. Heatmap of MSigDB hallmark pathway activities across CRC subclones; 1,000 cells were randomly sampled per subclone.

**E**. Primary versus metastasis prevalence of CRC subclones measured by *R_o/e_. R_o/e_* values were categorized as follows to reflect the degree of enrichment: +++, *R_o/e_* > 1; ++, 0.8 < *R_o/e_* ≤ 1; +, 0.2 ≤ *R_o/e_* ≤ 0.8; +/−, 0 < *R_o/e_* < 0.2; −, *R_o/e_* = 0.

**F**. Scatterplots showing Spearman correlation between the fraction of subclone 4 among malignant cells and the fraction of CD8+ T cells in the tumor microenvironment. Spearman’s correlation and *P*-value were shown.

**G**. Heatmap showing the proportion of subclones in malignant spots across 26 CRC metastatic samples.

**Figure 5**

**Comparison of CFMF with established methods.**

**A-B**. Boxplots of AUC values for genes selected by different methods across a range of number of genes (top 30-2,000) in PBMC3K dataset (**A**) and a representative GBM sample (**B**).

**C**. Boxplots of silhouette coefficients of different cell types calculated by different methods in PBMC3K dataset.

**D**. Boxplot illustrating SingleR annotation accuracy across cell types using genes identified by various methods in PBMC3K dataset across 50 iterations.

*P*-values were calculated by paired two-sided wilcoxon rank sum test. Significance: ***P* ≤ 0.01, ****P* ≤ 0.001, *****P* ≤ 0.0001.

**Figure S1**

**Validating CFMF with public scRNA-seq datasets.**

A. UMAP plots of PBMC3K dataset showing annotated cell types and canonical markers identified by CFMF.

B. UMAP plots of PBMC3K dataset showing novel markers identified by CFMF.

C. UMAP plots of the GBM dataset showing four GBM subclone signature scores and expression of the corresponding genes identified by CFMF.

**Figure S2**

**CFMF fuels the identification of rare cell types.**

**A-D.** UMAP plots of four representative normal lung samples showing annotation of ionocytes and neuroendocrine cells (left), seurat clusters (middle) and marker expression (right).

**Figure S3**

**CRC subclone detection and quality metrics.**

A. UMAP and violin plots of per-cell total unique molecular identifier counts (nCount) and number of detected features (nFeature).

B. Heatmap of Reactom pathway activities across CRC subclones. For each subclone, 1000 cells were randomly sampled.

**Figure S4**

**Molecular and cellular characteristics of CRC subclones.**

A. UMAP showing marker genes of distinct CRC subclones.

B. Dot plots of marker expression across CRC subclones.

C. Scatterplots showing the Spearman correlation between the proportion of each subclones among malignant cells and the proportion of CD8+ T cells in the microenvironment. Spearman’s correlations and *P*-values were shown.

**Figure S5**

**Comparison of CFMF with established methods in two GBM datasets.**

**A-B**. Boxplots of AUC values for genes identified by different signature gene selection methods with different number of genes (top 30-2000) in Abdelfattah2022 (**A**) and Wang2019 dataset (**B**).

**Figure S6**

**Comparison of CFMF with established methods in additional two GBM datasets.**

**A-B.** Boxplots of AUC values for genes identified by different signature gene selection methods with different number of genes (top 30-2000) in Darmanis2017 (**A**) and Neftel2019 dataset (**B**).

**Figure S7**

**CFMF improves tumor sample integration.**

**A**. UMAPs showing expression of *LGR5* and *TOP2A* when dimensionality reduction is performed using HVGs versus CFMF-selected genes.
