## Supplementary figures and images for "CFMF: A Clustering-Free Cell Marker Finder for Single-Cell Transcriptomics Data"

### main figure 1

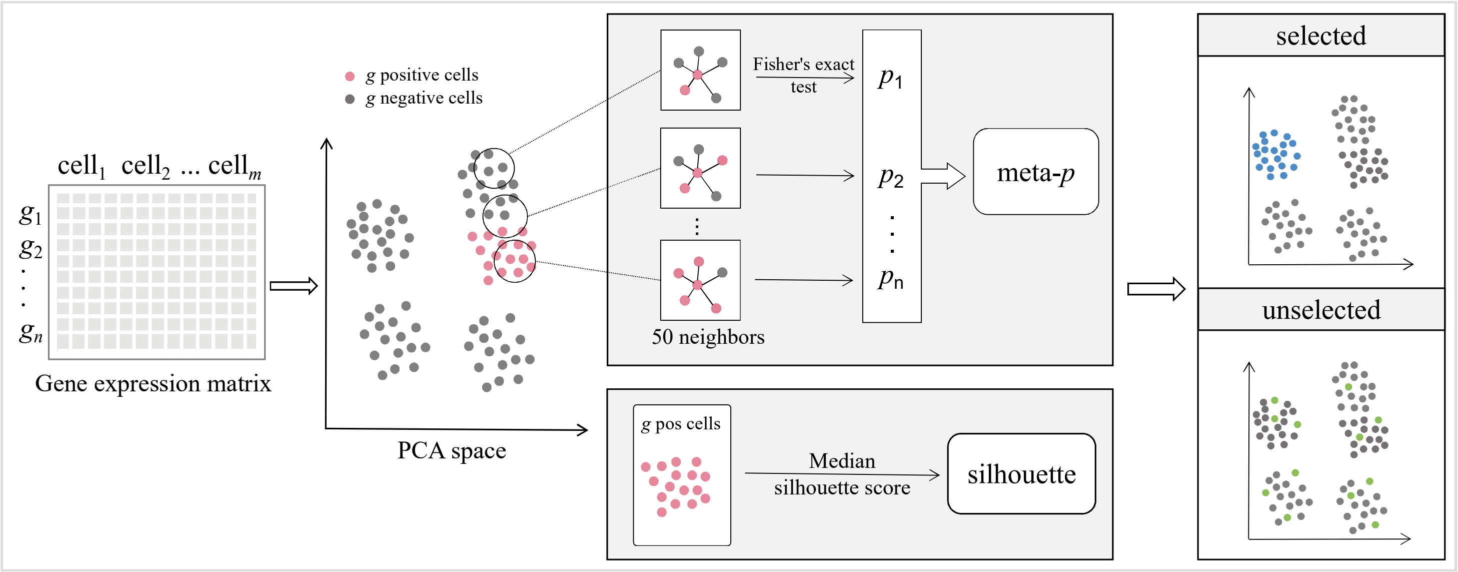

### main figure 2

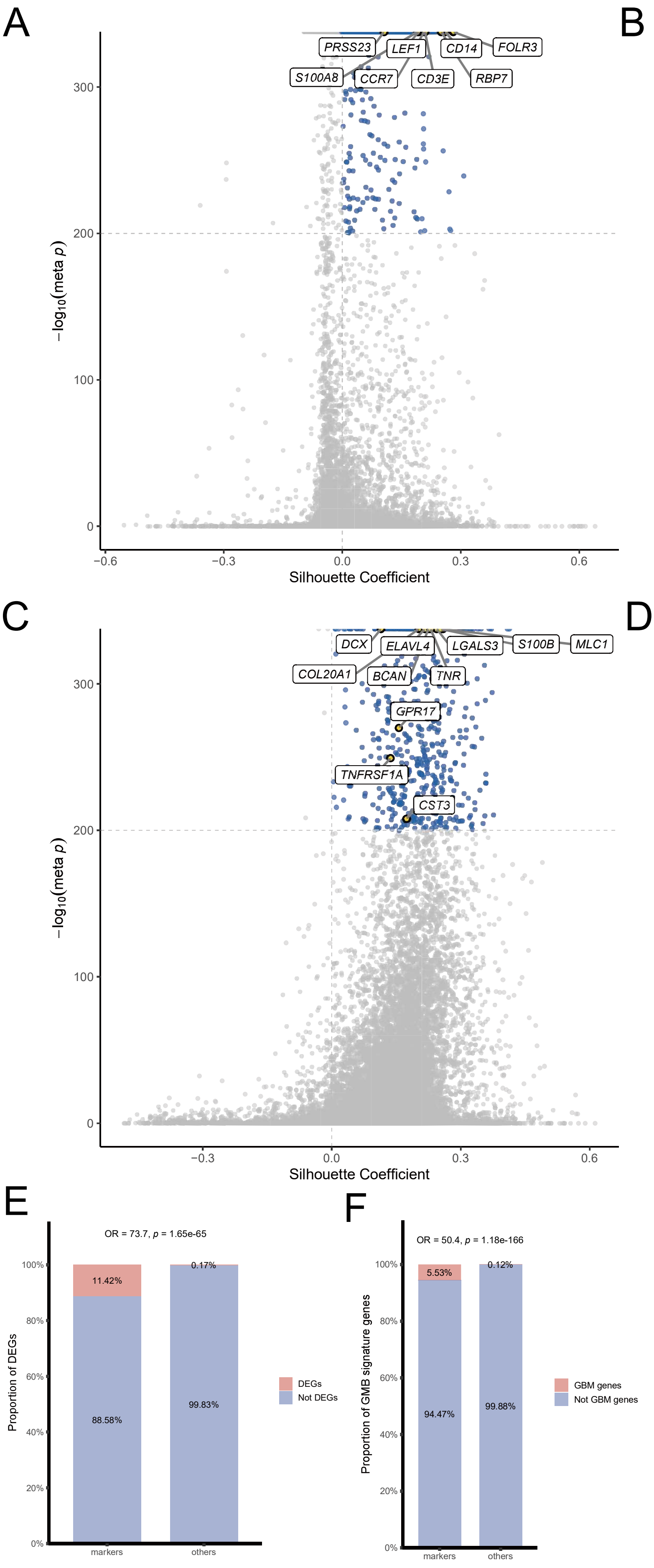

### main figure 3

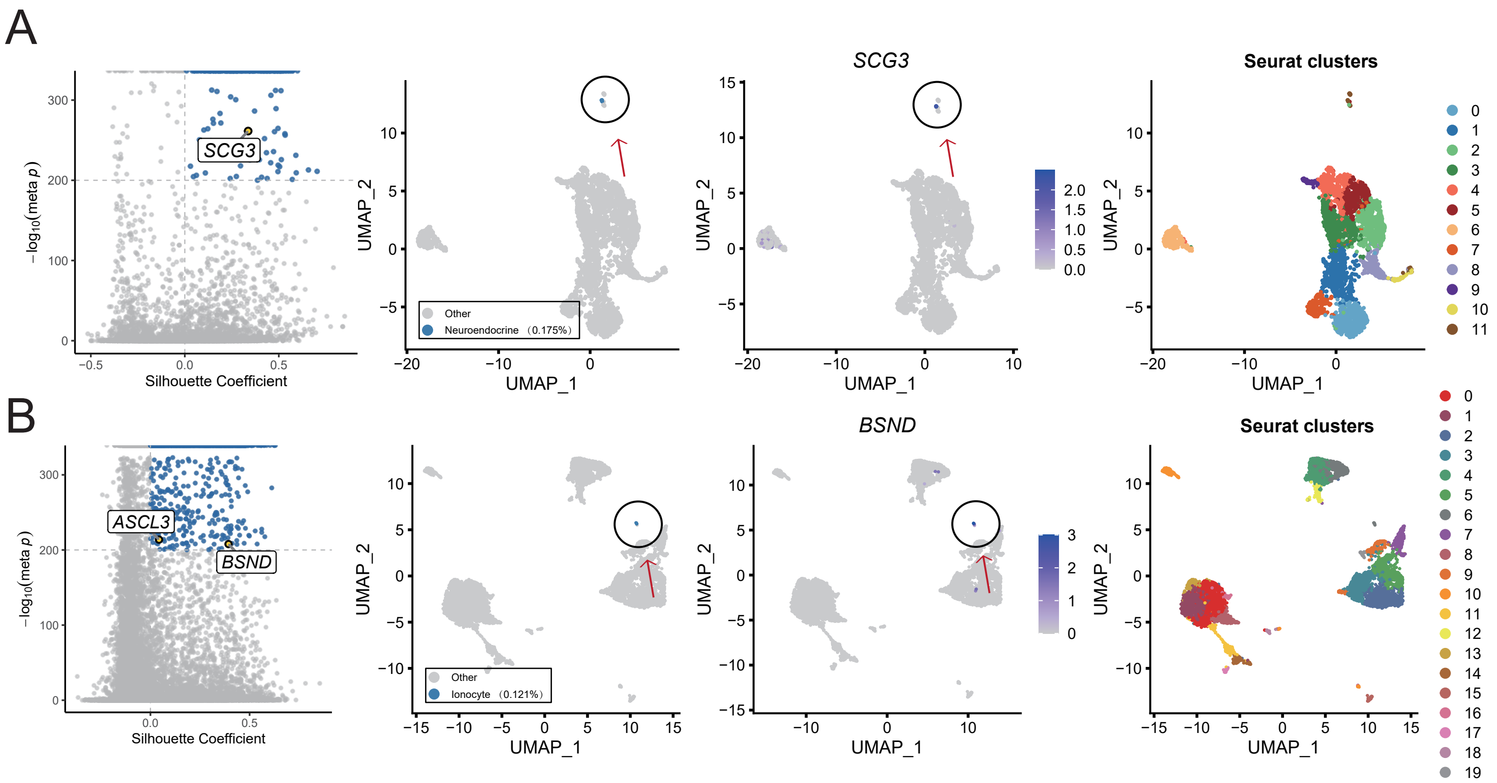

### main figure 4

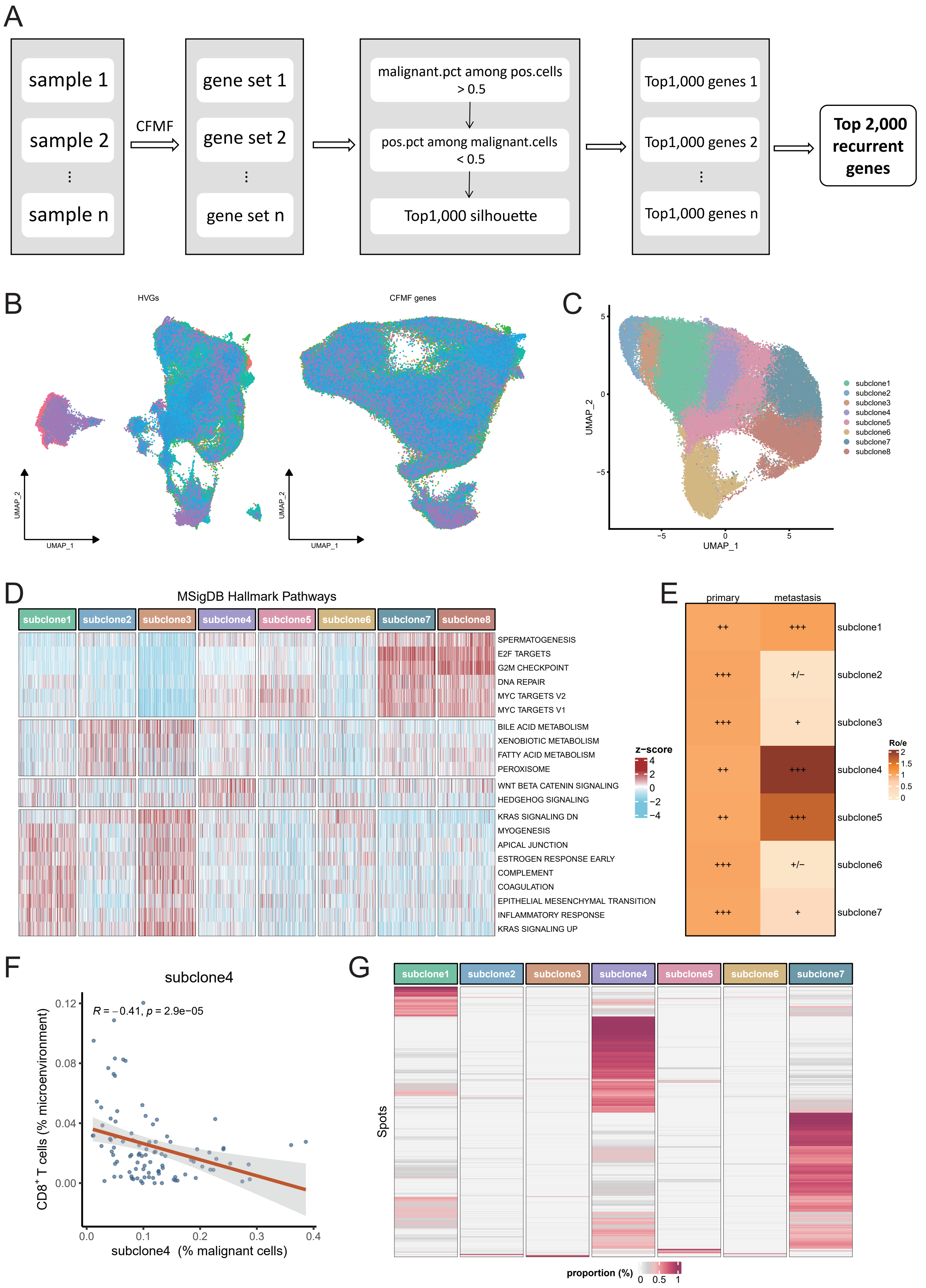

### main figure 5

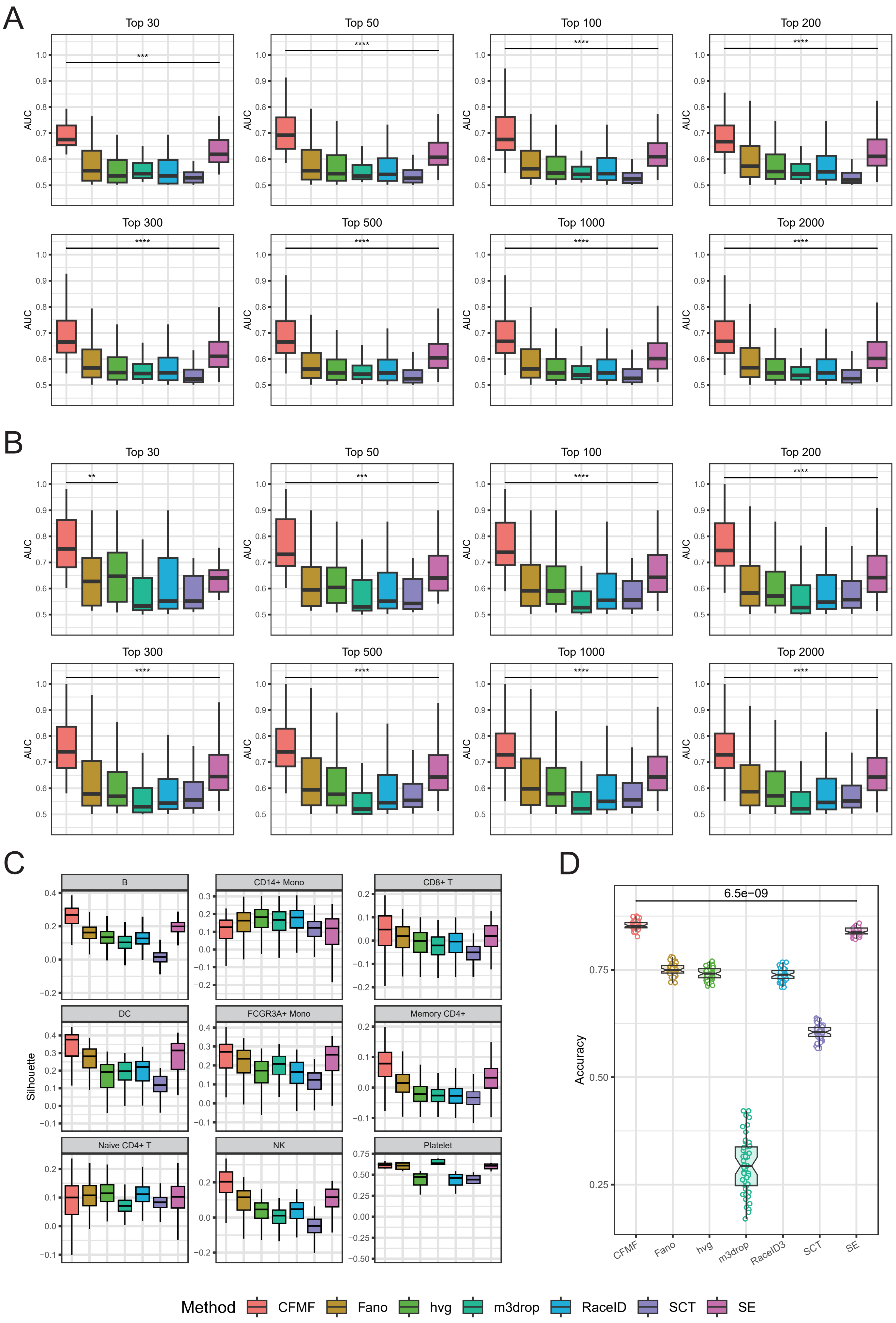

### supplemental figure 1

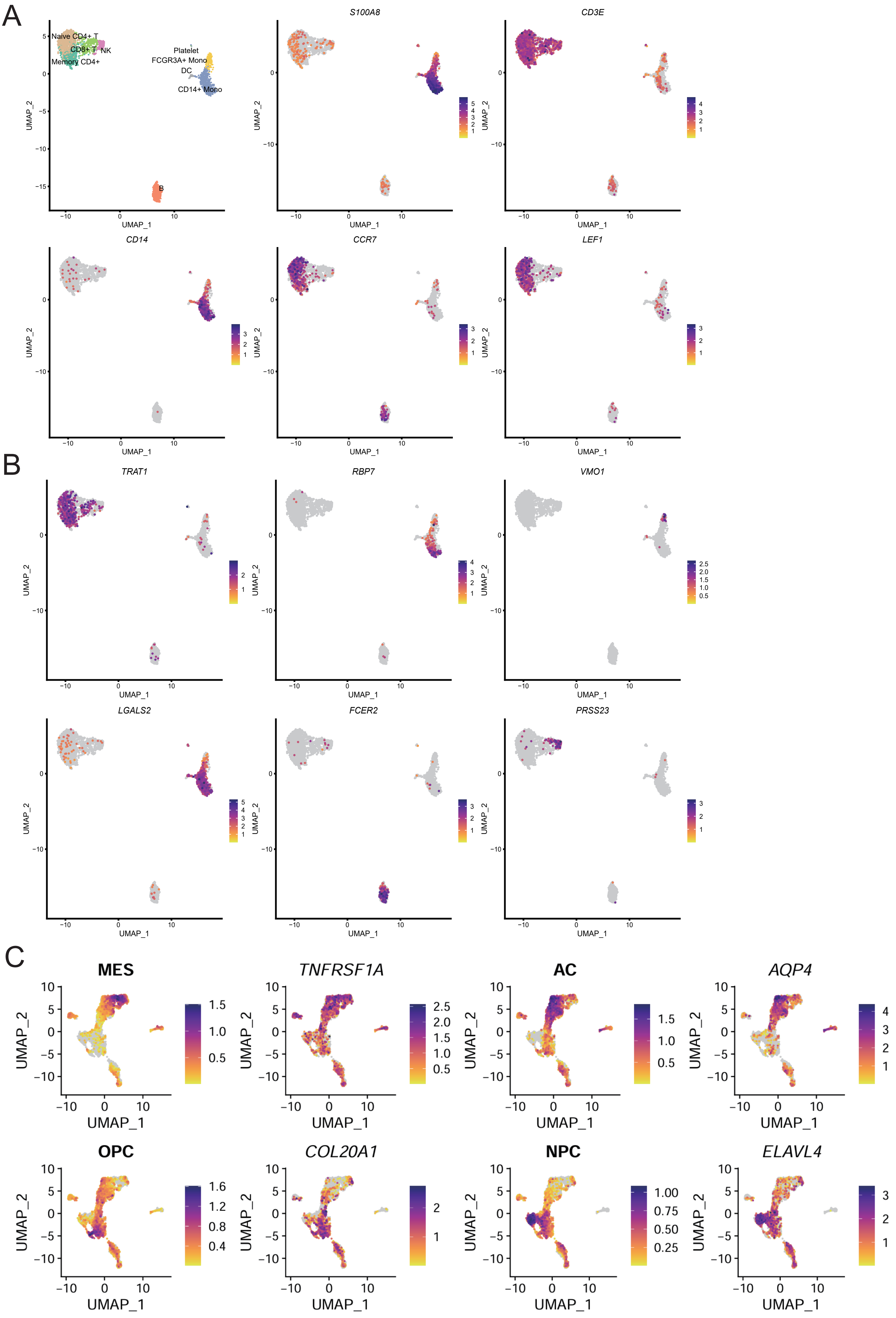

### supplemental figure 2

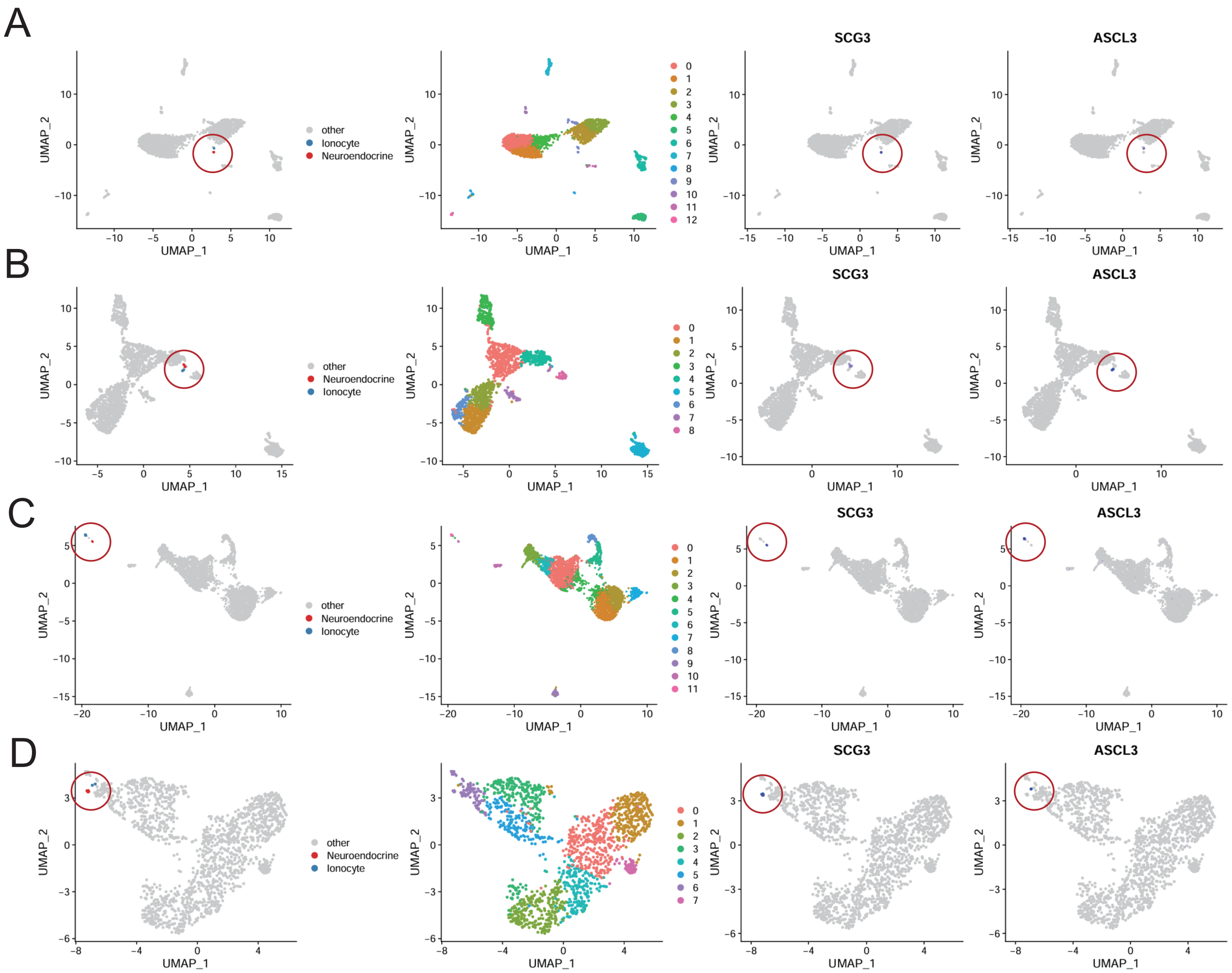

### supplemental figure 3

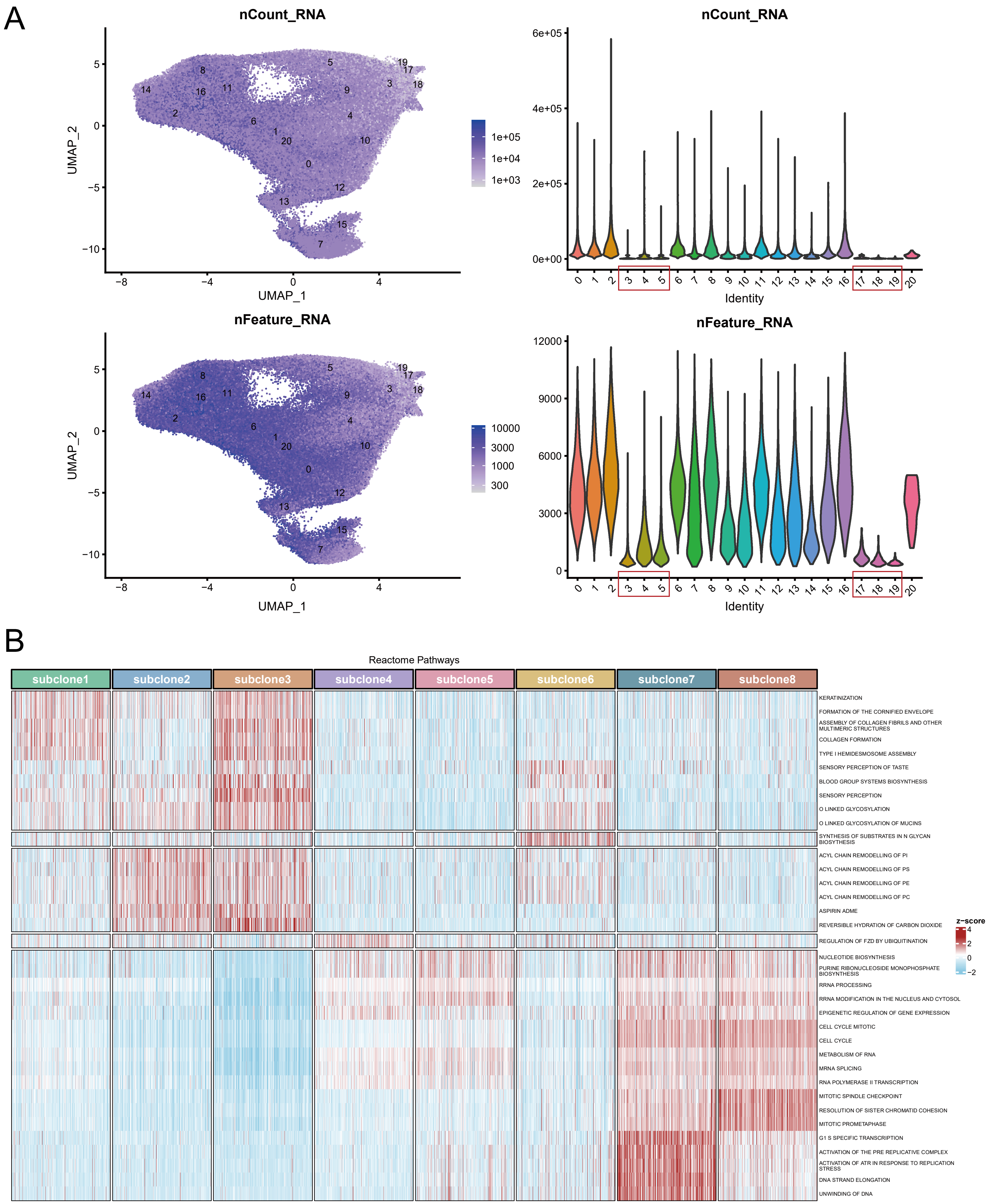

### supplemental figure 4

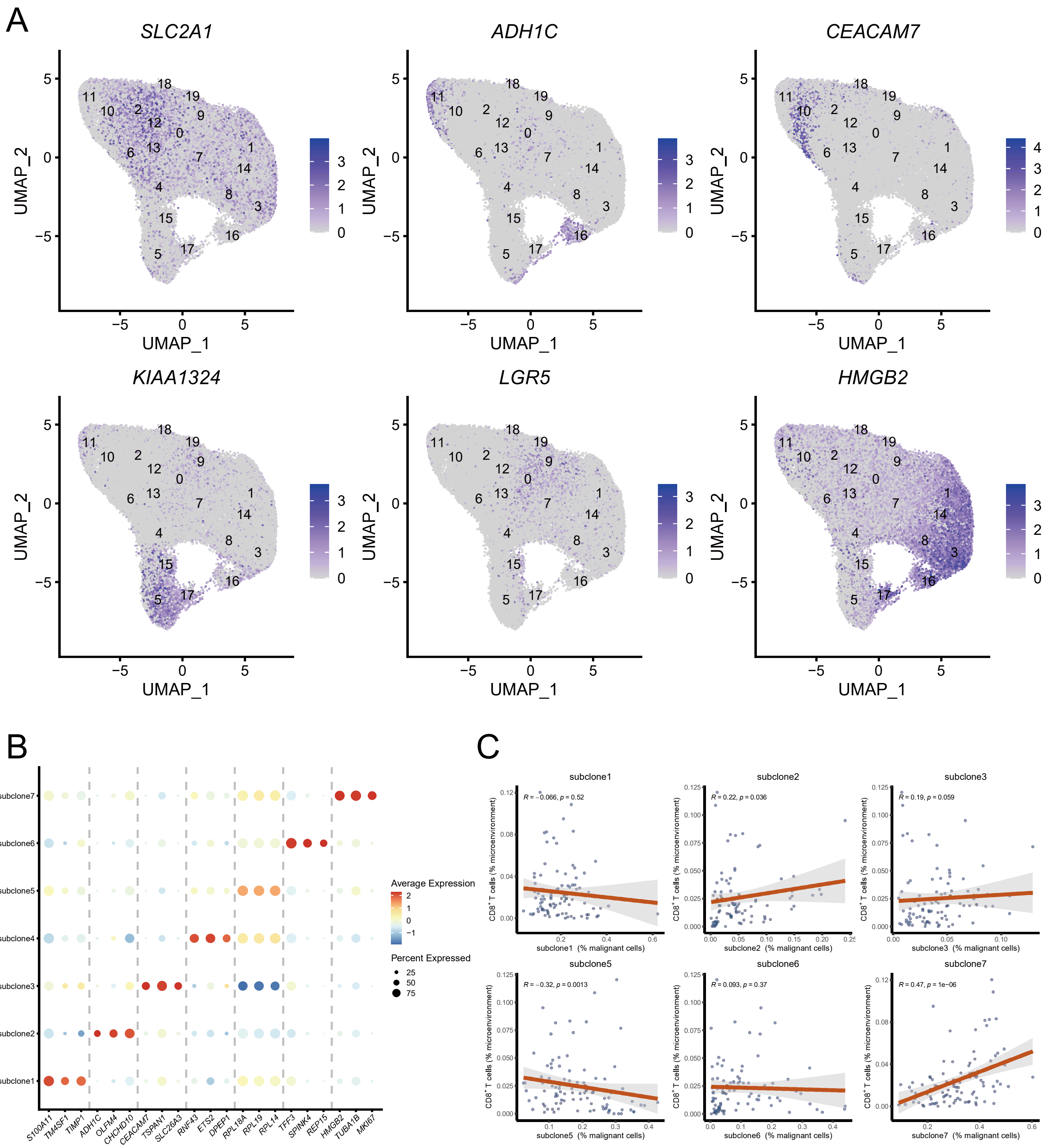

### supplemental figure 5

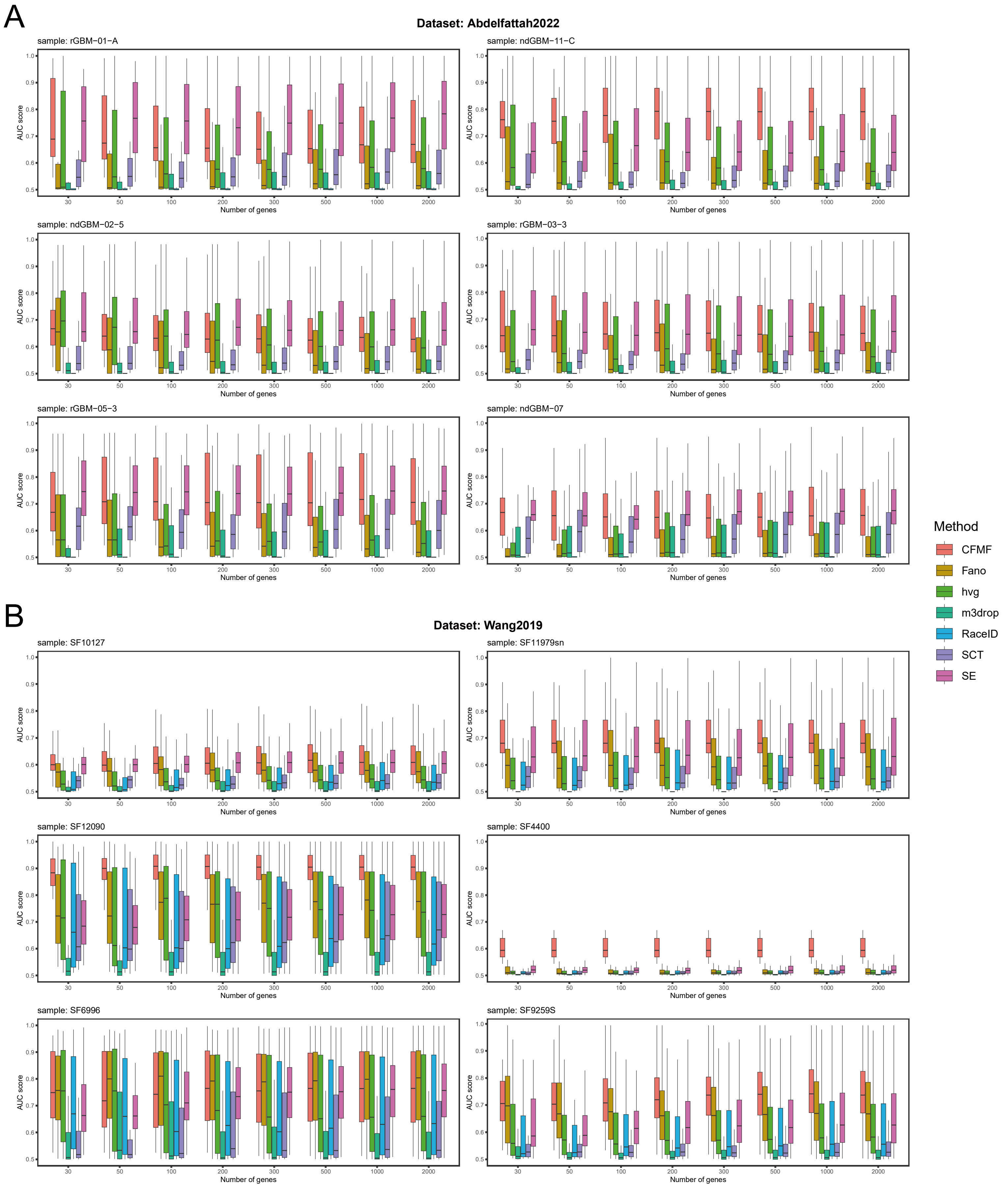

### supplemental figure 6

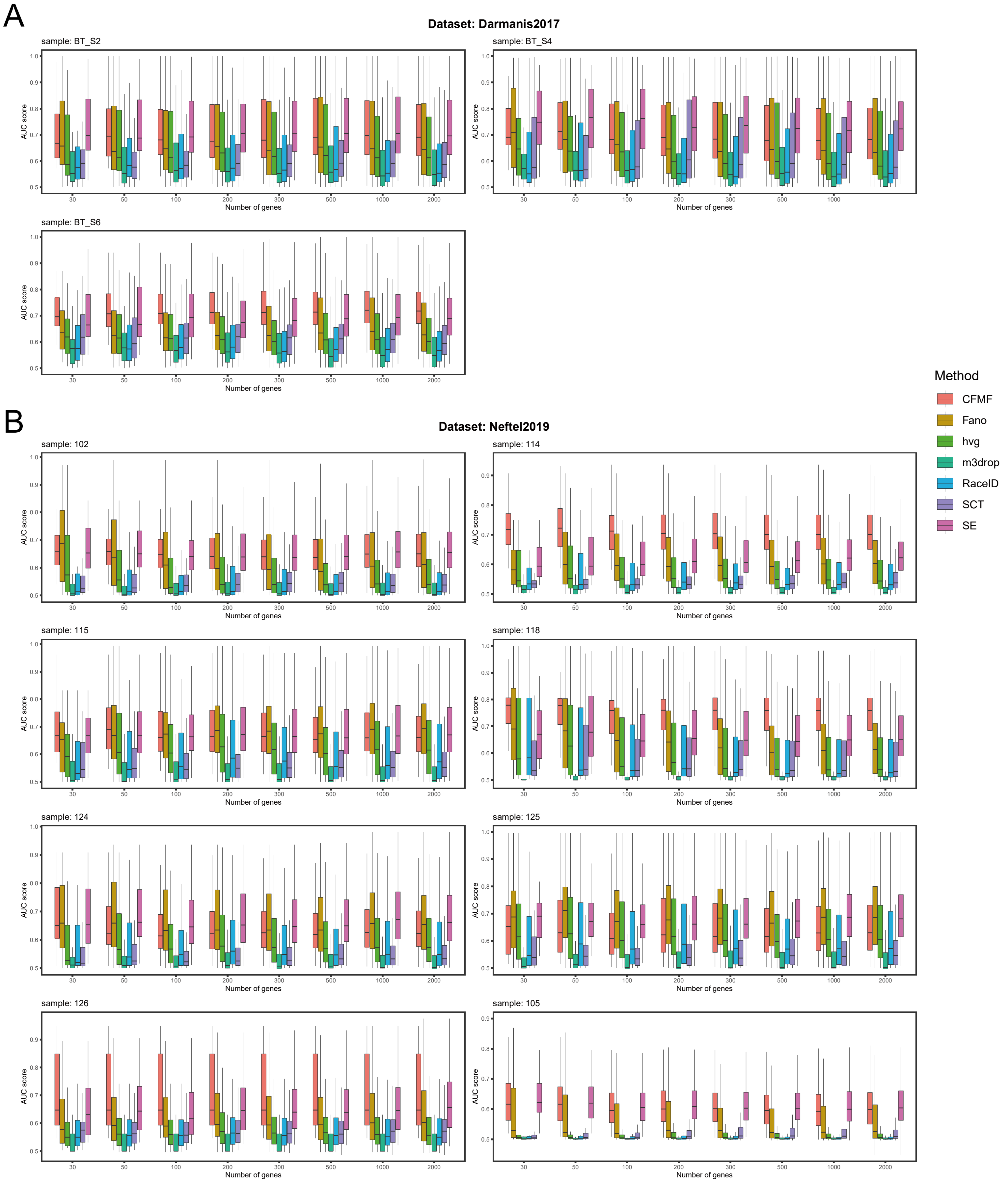

### supplemental figure 7

A

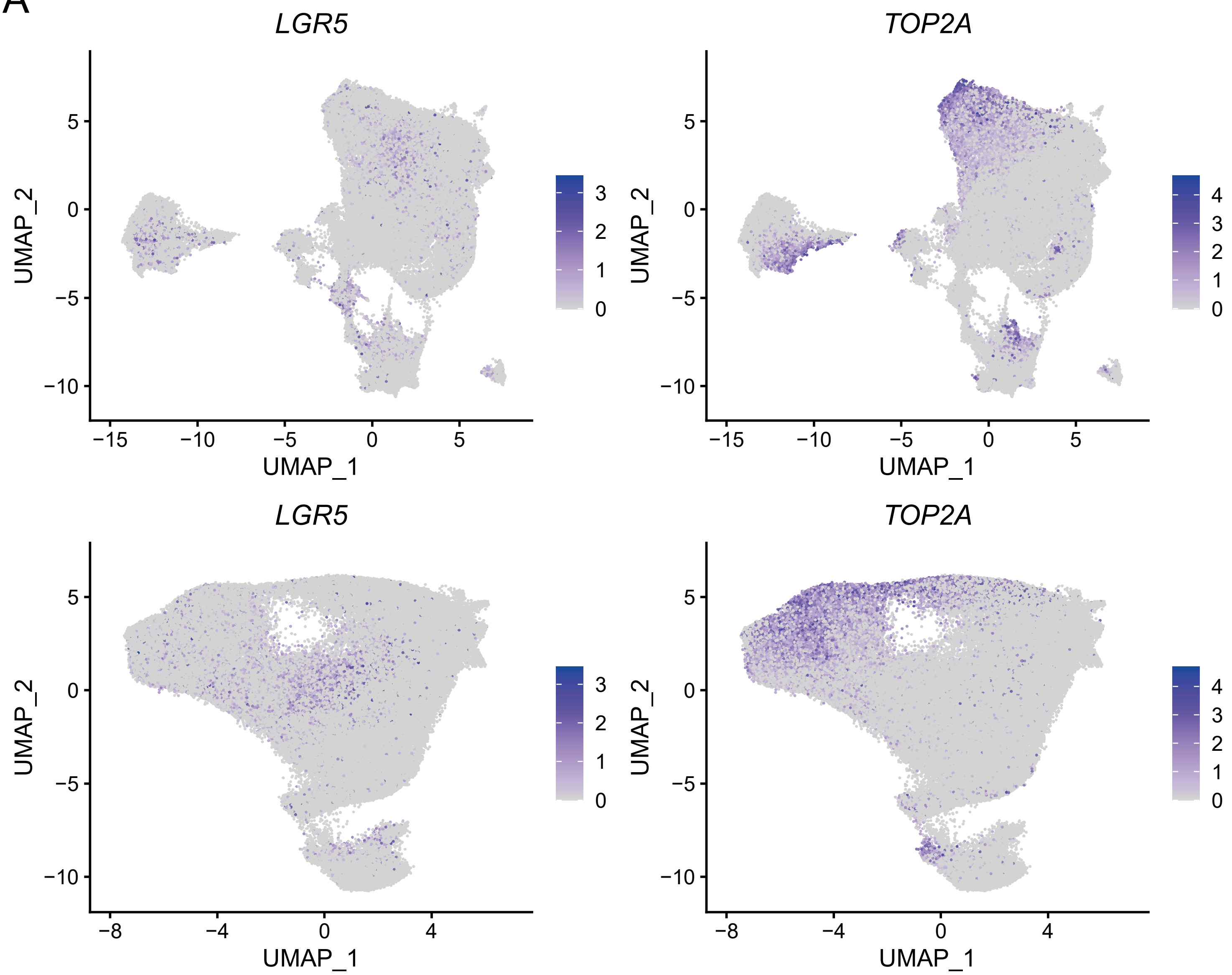
